# Mesoscale turbulence in the hippocampus

**DOI:** 10.1101/217877

**Authors:** A. Sheremet, Y. Qin, J.P. Kennedy, A.P. Maurer

**Affiliations:** Engineering School of Sustainable Infrastructure & Environment (ESSIE), 365 Weil Hall, University of Florida Gainesville FL 32611.; Department of Neuroscience, McKnight Brain Institute, College of Medicine, University of Florida, Gainesville, Florida 32610.; Department of Biomedical Engineering, University of Florida, Gainesville, Florida 32611

## Abstract

Wave turbulence provides a powerful stochastic description for nonlinear collective neural activity in the hippocampus. Recent studies that show theta waves propagating across the hippocampus suggest that turbulence is a natural description for collective neural activity. We formulate the fundamental principles of a turbulence model and demonstrate turbulent behavior by analyzing rat hippocampal LFP traces. LFP spectra and bispectra exhibit fundamental turbulent properties: weak nonlinear coupling, energy cascade, and stationary spectra of the Kolmogorov-Zakharov type (power-law). Weak turbulence holds the promise of quantitative physical models of hippocampal dynamics, in the service of understanding the mechanisms of multi-scale integration of brain activity to cognition.

---

The physics behind the observations of the scale-invariance of brain activity (power-law spectra *f*^*−α*^, where *f* is the frequency), has been subject of intense research and debate in recent years. Power-law spectra are associated with fractal geometry. For example, a simple recipe for a *f*^*−α*^ spectrum^1,2^ is to superpose pulses *X*_*T,τ*_ (*t*) centered at time T and with an exponential decay rate τ distributed as *F*(*τ*) ∝ *τ*^*−β*^. If the pulses are statistically uncorrelated (e.g., separated in time), the time series Σ_*T,τ*_ *X*_*T,τ*_ (*t*), has the power-law spectrum, with *α* = *β* − 2. The dilemma with this type of mathematics is that it does not address the physical source of scale invariance; it just shifts the “*f*^*−α*^” question to the equivalent “*τ ^−β^*” one. This is precisely the problem self-organized criticality^3^ hoped to resolve. Introduced as a universal physical model for scale invariance, it was quickly adopted as a possible brain model^4^, and is currently the leading large-scale representation of neural activity. However, after more than 20 years of intense research efforts, the theory has not progressed much beyond the sand-pile paradigm and simplistic numerical models, “many of which are not intended to model much except themselves”^5^. While it is not inconceivable that the brain is indeed in a self-organized critical state, the existing evidence is less than compelling. It seems that the search for a dynamical (as opposed to geometrical) description of the brain is still open.

An alternative dynamical formulation is suggested by recent studies^6,7^ that provide compelling evidence that the hippocampal theta rhythm is the expression of waves (wavelength ≈20 mm, phase speed ≈ 160 mm/s) propagating at various angles with respect to the dorsoventral axis. Wave fields can exhibit scale-invariant states described by power law wave spectra, whose nonlinear and stochastic properties are studied within the weak (wave) turbulence formalism. Although the basics of the turbulence formalism are rather straightforward (see Methods), to our knowledge, no attempt has been made so far to examine the physics of brain activity from the wave-turbulence perspective.

Originally formulated for hydrodynamics, the scope of turbulence has expanded through the work of Richardson, Kolmogorov, and Zakharov ^8–10^ to become the theoretical foundation of physical disciplines ranging from plasma physics, to nonlinear optics, Bose-Einstein condensation, water waves, coagulation-fragmentation processes, and many others^9,11,12^. The unifying element in this diversity is the type of physical system studied. A “turbulent system” satisfies two essential requirements: 1) it has a large number of components, that 2) interact nonlinearly. The “large number” requirement implies that the physics governing the components, and their characteristic spatial and temporal scales are distinct from the system. We will refer to the system scale as the macroscale, and to the component scale as the microscale. The second requirement, nonlinear interaction, describes the physics of the components, and plays the crucial role of connecting evolution across scales. If energy is introduced into the system at a given scale, nonlinear interactions will spread it to nearby scales. This flux of energy across scales is called the *turbulent cascade*^8,10^, and is the fundamental property defining turbulence (Fig. 1a).

**Figure 1:**
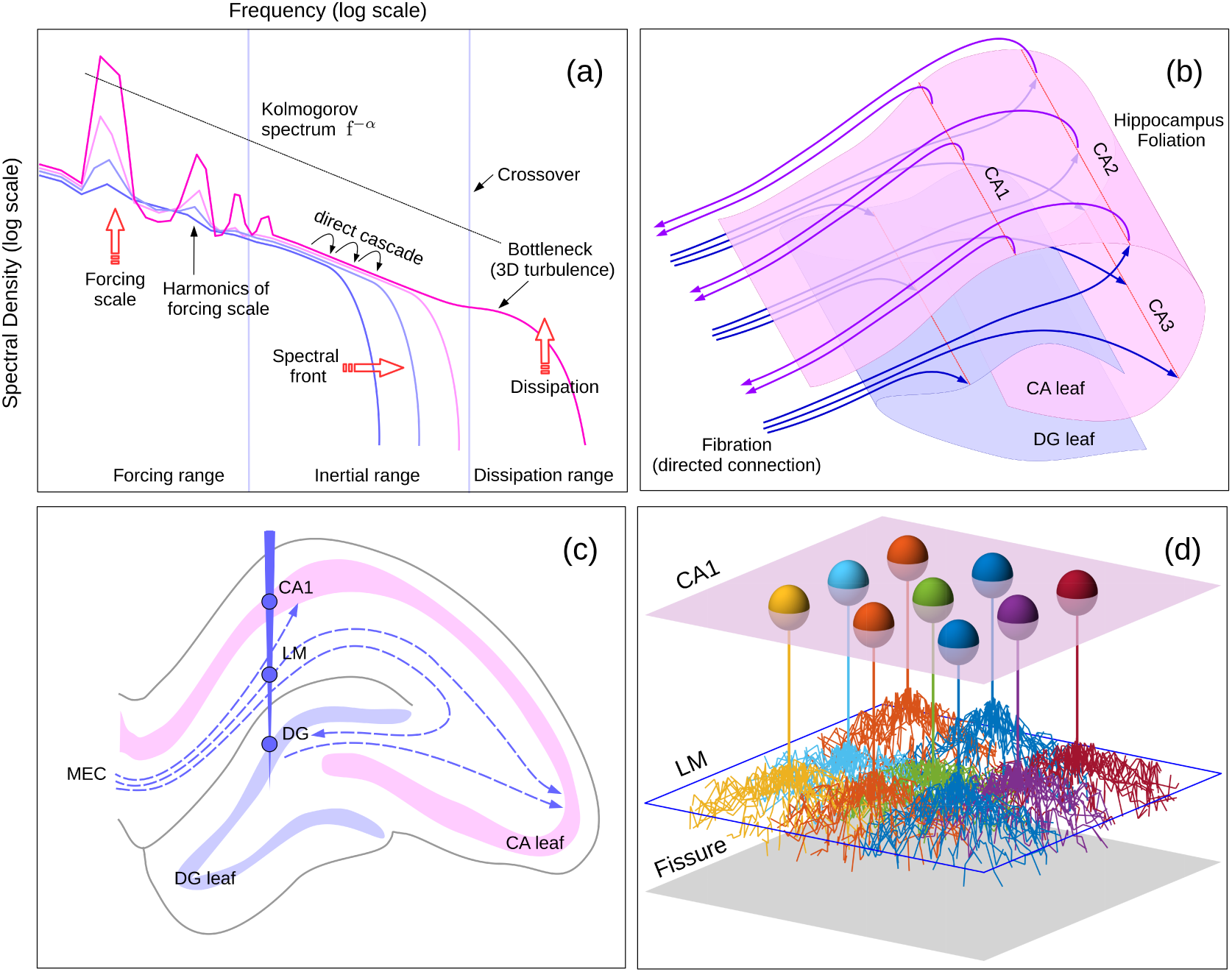
a) A schematic of a possible evolution of a hydrodynamic turbulence spectrum^11, 37^. Frequencies represent temporal scales. The dissipation range is characterized by a spectral decay faster than any power law. In the inertial domain, energy flows across scales from the forcing range toward the dissipation range. This energy flux is called turbulent cascade. As energy is pumped into the system, a spectral front develops that moves toward higher frequencies, leaving a stationary KZ spectrum in its wake (dashed line indicates the slope). When the spectral front crosses the molecular dissipation crossover, a small bump occurs in three-dimensional turbulence (crossover bottleneck^38^). b) A simplified schematic of the hippocampus fibration-foliation complex. A foliation is a collection of two-dimensional “leaves”, lattices of neurons (folded surfaces). Because the fibration-foliation distinction serves modeling purposes, the boundaries of the leaves are defined by the functionality studied (for example CA could be subdivided into CA1-2-3). Fiber bundles (arrows) are directed connections performing input/output functions. c) Schematic of rat hippocampus with the location of LFP sites analyzed in this study. d) Schematic of the CA1/LM structure, represented as plane lattices. The CA1 lattice (pink plane) is the locus of the somas of pyramidal cells (colored spheres). The LM region (blue outline) circumscribes the synaptic connections of the CA1 cells. The fissure is represented as gray plane.

Brain activity spans a spectrum of scales. A persistent challenge posed by higher-cognition is understanding how these scales integrate into global brain coordination. At cell scale, action potentials (frequency*≈*10^3^ Hz) provide the “atomic” constituents of activity. At global-scale, theta, with a frequency three orders of magnitude lower, is believed to provide a temporal structure around which smaller scale oscillations organize^13^. Neither of these oscillations can represent cognition. The first step in understanding integration is understanding how the dynamics of intermediate scales connect to large scale activity. Hippocampal activity is a paradigm for this problem^13^. Turbulence provides a powerful approach that might explain its intriguing features, such as the weak phase coupling between theta and gamma rhythm, which seems to speak to scale integration; and power-law shapes *f*^*−α*^ with multiple slopes *α* depending on the recording site, seemingly stretching the scope of criticality theory. Below, discuss the elements of a weak-turbulence model for the mesoscale brain, examine evidence of turbulence in brain activity, and discuss possible implications of such a model in understanding some of the more intriguing physical features of brain activity.

## Anatomy of mesoscale turbulence

If the brain is simply a bundle of neural *fibers* used to convey non-overlapping, non-interacting pulses of information, turbulence does not seem applicable, as randomness and nonlinearity interfere with this function. However, representing the brain as a collection of neural fibrations is only an approximation. The typical connectome^14^ voxel is itself macroscopic (*>* 10^6^ neurons or more^15^). At smaller scales, the topology changes. Observations of wave propagation in the hippocampus^6,7,16^ suggest that, local connections are favored over distal ones^17,18^, and that parts of the hippocampus, e.g., cornu ammonis (CA), and the dentate gyrus (DG) regions function as a collection of two-dimensional neural lattices (topologically, a *foliation*; e.g., ^19^). The CA1 leaf is connected to the rest of the brain through the fibrations in the radiatum and lacunosum moleculare (LM) region: the Schaffer collaterals, perforant pathway and hippocampal commissure. Fibrations dominate the macroscale^14,15^, defining the large-scale hierarchical structure of the brain and global physiology. So far, the data suggests that foliations occupy an intermediate (meso) scale between macroscale and the microscopic, cellular constituents.

If neural activity propagates as waves on foliations, overlapping in space and time becomes possible. Therefore, nonlinearity and stochasticity become relevant. Mesoscale collective activity is stochastic (the state of a single neuron is inconsequential) and inherits the intrinsic nonlinearity of individual neurons. However, the nonlinearity has to be weak (e.g., strong nonlinearity is a symptom of pathology^20^), implying that interacting waves never develop strong phase correlations, and remain statistically independent in the leading order.

Based on these considerations, we postulate that the brain activity on foliations can be represented as a multiple-scale, weakly-nonlinear wave field, whose dynamics is governed by turbulence. We shall refer to this as “mesoscale turbulence”. The wave field is the “large system”; the components are “elementary waves”, e.g., in a mathematical formalism, orthogonal Fourier modes. The dispersion relation defines the connection between temporal and spatial scales^21^, and the linear properties of the system: phase and group speeds, and conservation of energy. Weak nonlinearity suggests that *mesoscale turbulence is weak* (an example is the system of waves propagating on the surface of a deep ocean^22,23^). If this is true, standard averaging methods can be applied to the dynamical equations to determine the macroscopic behavior of the system^9,24^. In contrast to strong turbulence, weak turbulence allows for the formulation of universal equations for the evolution of turbulent systems^9,11,23^.

## Physiology of mesoscale turbulence

Here, we examine the spectra of LFP traces recorded from rats trained to traverse a circular track for food reward (see Methods, and Supplementary material). Although the functional form of the dispersion relation is not available, to translate frequency information into spatial scales, the minimal assumption that the relation is monotonic^**?**^ will suffice. This simply means that lower LFP frequencies correspond spatially to longer waves, thus engaging larger populations of neurons.

Figures 2-4 show estimates of LFP spectrum and bispectrum in three regions of the rat hippocampus: CA1, LM, and DG (Fig. 1). Layer localization was conducted through current source density analyses (see Supplemental Methods). To capture spectral evolution at incipient rat activity, and possibly the initiation of the turbulence cascade, the spectra were chosen to represent time intervals with low LFP variance. The LFP variance is related by a one-to-one mapping to rat speed3b, which is a direct measure of of behavior.

**Figure 2:**
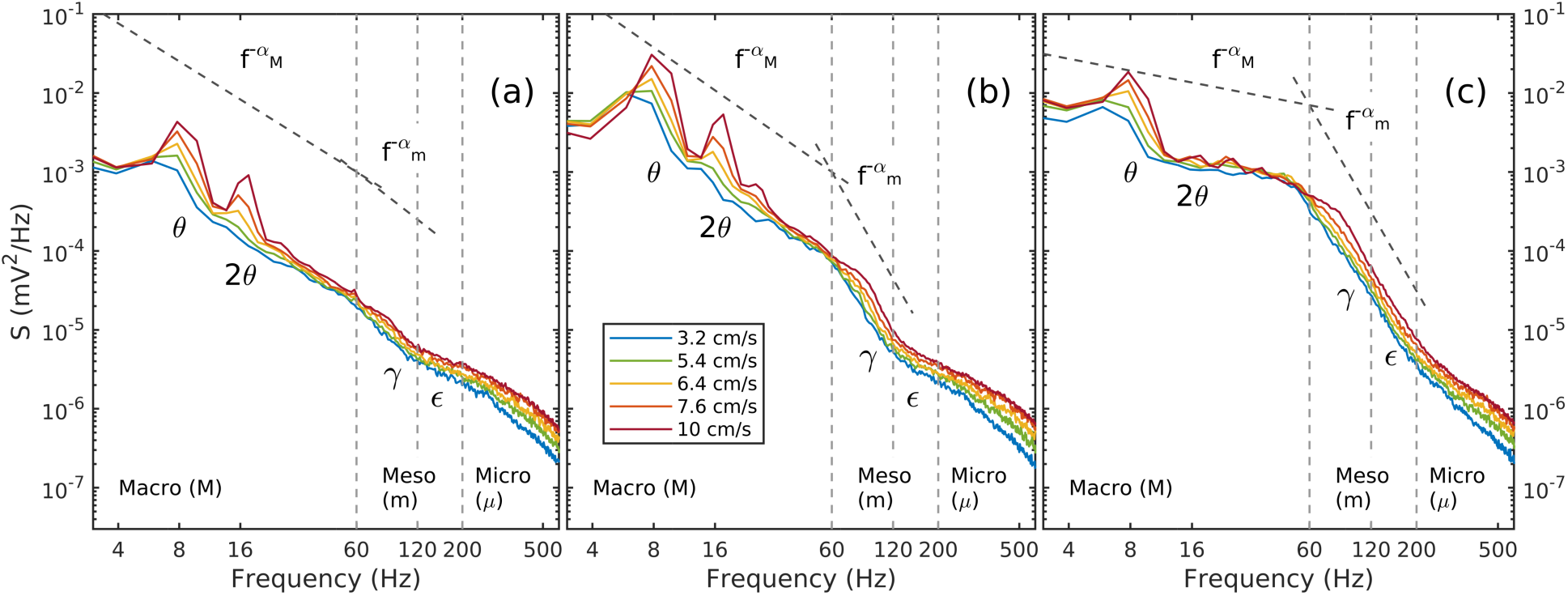
Evolution of the hippocampal LFP spectra for four velocity intervals, for regions a) CA1, b) LM, and c) DG. Three scales can be identified by the distinct slopes of the power law (*f*^*−α*^) spectrum: macroscopic, 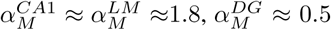, mesoscopic, 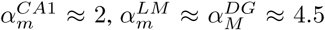, and microscopic, with decay faster than a power law. Crossover points, associated with a change in spectral slope (change in governing physics), are indicated by vertical gray dashed lines. Spectra correspond to the speeds marked by circles in Figure 3b.

The spectra in Figure 2-3a exhibit a number of obvious features: a developing peak at 8 Hz (theta), *f*^*−α*^ shapes, with two distinct slopes that vary depending on the hippocampus region (Buzsaki and Draguhn^21^ describe a single-slope spectrum), as well as a spectral tail decaying faster than a power law. The points points where the spectral slope changes (Fig. 2), called crossover points, indicate a change in the geometry (self-similarity) of the time series^25^. Crossover points clearly distinguish between three frequency intervals:

**Figure 3:**
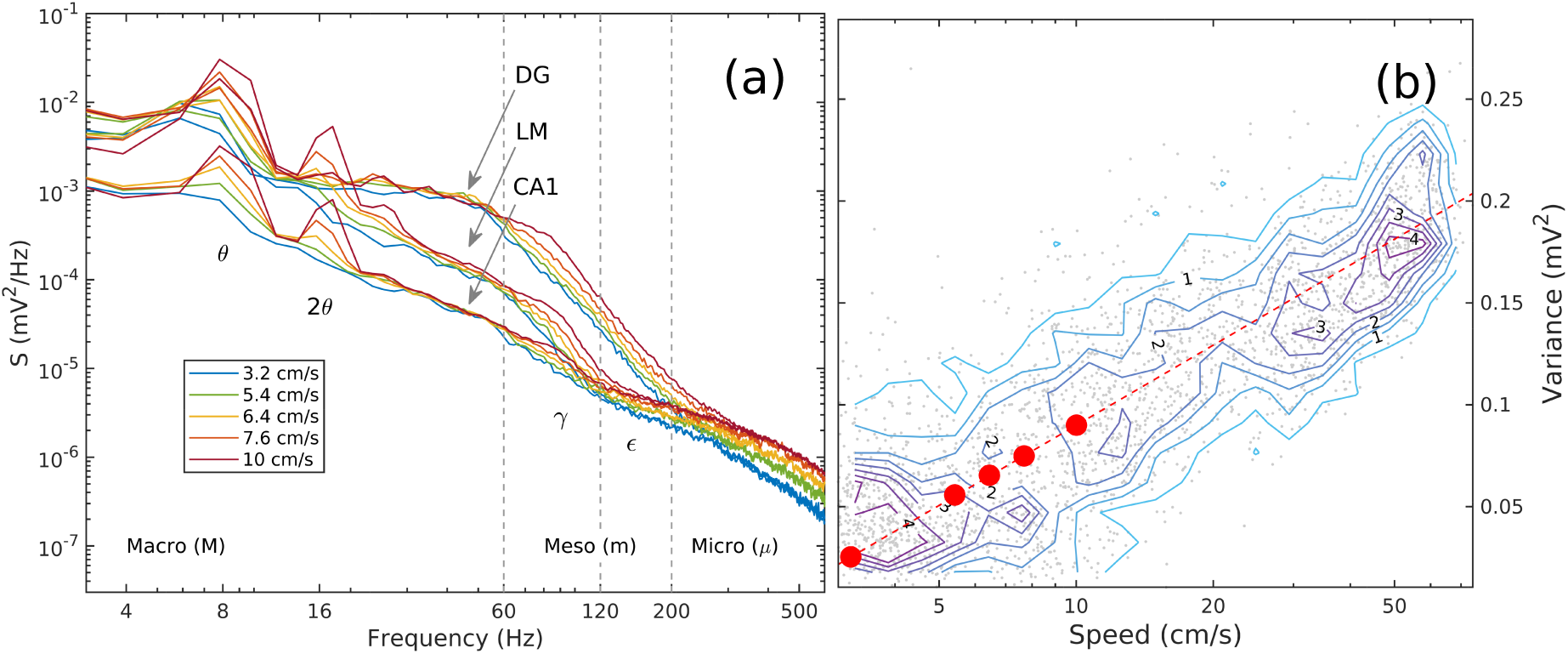
a) Superposed hippocampal LFP spectra recorded in CA1, LM, and DG regions (same as Fig. 2), highlighting the similarities and difference of the three regions. Spectra overlap in the dissipation (microscale) range, and the front-like KZ spectra in the inertial (mesoscale, gamma-epsilon) range can be seem clearly. b) Joint probability density function for LFP variance and rat speed. The dashed line is a best-fit analytical expression *v* = *a* log_10_ *V* + *b*, with *a* = 2.2 and *b* = 1.6. Circles mark the values of variance for which the spectral densities were estimated, midpoints of intervals 0, 0.05, 0.06, 0.07, 0.08, 0.09, and 0.1 mV^2^, with corresponding speeds are approximately 3.2, 5.4, 6.4, 7.6, and 10 cm/s. The relation between variance and speed follows the Weber-Fechner law of stimulus perception^39, 40^

Macroscale (*f* < 60 Hz). This frequency band is dominated by the theta rhythm and its harmonics^26^. The low frequency suggests that it is associated with macroscale (subscript *M*) processes, possibly mostly with fibrations. The spectrum follows a power-law, with a slope that changes by region: LM and CA1 have approximately the same slope 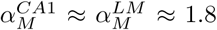; the DG spectrum is much flatter,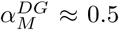. We avoid using the “slow gamma”, traditionally ascribed to the 20-50 Hz or 30-80 Hz bands^27,28^, because our data did not allow for the disambiguation of the theta harmonics with slow gamma.

Mesoscale (60 < *f* < 200 Hz). This is the domain of the gamma and epsilon rhythms (subscript *m*; usually associated with the 60-90 Hz and 120-200 Hz, respectively). The name epsilon is preferred to “fast gamma” in congruence with previous publications^29, 30^ which suggest that gamma is generated through local volleys of activity of coupled interneuron-interneuron (I-I), or pyramidal-interneuron (E-I) cell pairs, while epsilon results from spike-afterdepolarization and spike-hyperpolarization components. The spectrum follows a power law, but the structure is complicated by the different behavior of the two sub-bands. Near CA1 and LM, gamma (60 ≤ *f* ≤ 120 Hz) dominates, with low-power epsilon oscillations; 120 ≤ *f* ≤ 200 Hz in the DG region, covering both gamma and epsilon. The spectral slope is the same in LM and DG regions, 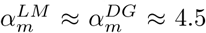, but much smaller in CA1, where 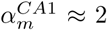 is also closer to the macroscale value 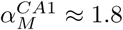. Spectral and bispectral analysis (see below) define the gamma and epsilon bands in a natural way, free of a priori assumptions.

Microscale. This is the high-frequency domain (subscript *µ*), where spectral decay is faster than a power law. The dominant processes include action potentials, fast synaptic time constants, ion channel opening and closing and, at very high-frequencies, heat dissipation. Near CA1 and LM, the lower limit of the microscale range extends over the epsilon band *f* > 120 Hz; near DG, where epsilon has higher power, the lower limit is *f* > 200 Hz.

It is worthwhile recalling that these spectra are observed on a foliation, where we conjecture that all oscillations are represented by waves, even if some (e.g., macroscale oscillations, theta) have a different nature in other parts of the brain. In the hippocampus, all oscillations become connected through the nonlinearity of the supporting neural lattice, and the spectra are expected to reflect their underlying nonlinear and stochastic dynamics.

## Physics of mesoscale turbulence

Our conjecture of weak mesoscale turbulence implies that 1) a kinetic description can be derived for the spectral density of energy, and 2) stationary Kolmogorov-Zakharov^11^ (KZ) spectra of the form *f*^*−α*^ are possible. We expect that voluntary rat activity should be associated with statistically stationary brain states, by virtue of being sustainable for an indeterminate period of time. It follows that KZ spectra are realized and can be observed in the hippocampus. The spectral slope *α* of a KZ spectrum is important because it characterizes the dimension and the dynamics of the system (see Supplementary Material; with the slope being indicative of cascade strength). The frequencies where the spectral slope changes (Fig. 2) are called crossover points, and signal a shift in the dominant physical processes.

As expressions of mesoscale turbulence, the spectra in Figure 2-3a provide information about the dynamics of collective neuronal activity. A comparison with typical turbulence (Fig. 1a and 2) shows compelling similarities and also interesting differences. The similarity between theta nonlinearity (evidenced by harmonic excitation^26^) in the macroscale domain and the forcing range in Figure 1a suggests that theta plays the role of a spectral window for energy input into the hippocampus. Like the dissipation domain in Figure 1a, the microscale domain *f* > *f*_*mµ*_ exhibits a faster than power-law decay, as well as a bottleneck bump (see Fig. 1a). If the forcing and dissipation scales are well separated, there exists an intermediate range, the Kolmogorov “inertial range”, where nonlinear interactions allow energy to cascade toward the dissipation range. *This suggests that the mesoscale is inertial*. Our equilibrium conjecture identifies the mesoscale range as a KZ stationary spectrum, with *constant spectral flux* (energy) cascade.

The features that set the hippocampal spectra apart from the simple turbulence schematic in Figure 1a speak to the complexity of the brain. The multiple-slope spectra reflect the distinct physics governing the different scales of activity. Also, pure spectral fronts as shown in Figure 1a exists only when a system starts from rest, i.e., with zero energy in the inertial range. But the brain is never at rest in this sense of the word. The hippocampus, as a wave-supporting lattice, is globally connected and channels some of the global brain energy. Therefore, hippocampal spectra contain a extraneous background level of activity that is never zero.

The variability of the spectral shape within the hippocampus is striking. Overall, the CA1 spectrum responds the least to behavior, and preserves an nearly-constant slope through the macro- and mesoscale ranges. In the LM and DG regions, the mesoscale (gamma) range responds strongly to the initiation of activity. As rat activity increases, the theta slope begins to extend into the lower-frequency end of the mesoscale domain, “pushing” energy into the gamma range, increasing its variance and forming a secondary KZ spectrum that propagates like a front toward higher frequencies. Throughout this evolution the spectral slope remains constant *α*_*m*_ ≈ 4.5, consistent with the stationarity hypothesis, and suggesting similar turbulent dynamics between DG and LM. By comparison with the DG observations, gamma waves are weaker in LM, and epsilon waves are low-power, indistinguishable from the dissipative range.

The penetration of the macroscale spectrum into the the lower gamma range, and the development of the mesoscale KZ spectrum are strongly suggestive of a cascade process. The bispectra support this assumption. Low-speed bispectra (Fig. 4) show barely-significant phase coupling: CA1 is nearly-linear; weak coupling can be seen between theta and its second harmonic (LM and DG); and theta-gamma coupling (DG). A the higher running speeds (i.e., >40 cm/s; large variance, or strong theta forcing, Fig. 5), bispectra show a well defined increase in theta nonlinearity^26^in all three regions (up to 4 harmonics are easily identified). The nonlinear structure of the CA1 LFP remains the simplest, while in the LM and DG regions the phase-coupling landscape becomes progressively richer, shedding light on the origin of the structure of the mesoscale KZ spectra. In the LM, despite the weak epsilon presence, the bispectra show unequivocally a strong phase coupling of both theta-gamma and theta-epsilon (Fig. 4; see also the schematic in Fig. 6b). Indeed, these observations suggest that theta exerts a nonlinear forcing of gamma waves in the LM, pushing their variance up and establishing a KZ spectrum that grows as rat activity increases. Epsilon oscillations in the LM, while being low power, exhibit coupling to theta as well. The reason for the coupling is that the epsilon rhythm is a consequence of spike-afterdepolarization and spike-hyperpolarization components^29,31^. This is consistent with high frequency activity, such as back-propagating action potentials into the dendrites of the LM, being phase coupled. In contrast, high-speed bispectrum near the DG region shows no theta-epsilon coupling, but shows a distinct gamma-epsilon coupling of the type related to second harmonic generation. Stated differently, the origin of epsilon activity in the LM is relayed spiking activity from pyramidal neurons in CA1. In the DG region, however, epsilon frequencies are the result of energy transfers nonlinearly interacting populations of neurons at different scales, generating harmonics in a manner similar to theta harmonic generation (see Discussion).

**Figure 4:**
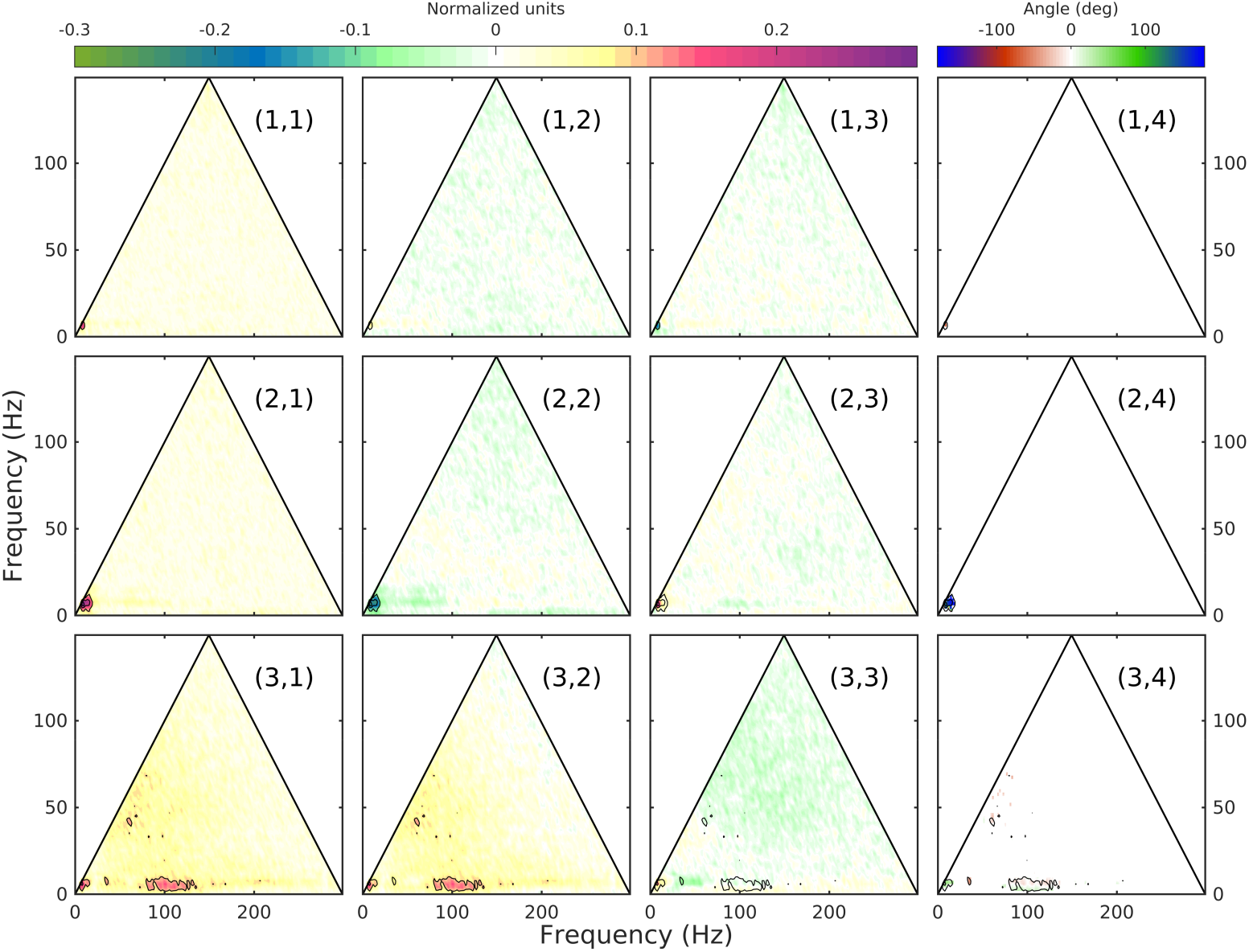
Bispectra of LFP recordings corresponding to low rat speed, *v* < 10 cm/s. Columns: 1) bicoherence (normalized modulus of bispectrum); 2) skewness (normalized real part of bispectrum); 3) asymmetry (normalized imaginary part of bispectrum); 4) biphase (argument of bispectrum). Rows: 1) CA1; 2) LM; 3) DG. See Discussion and Figure 6 for an interpretation.

**Figure 5:**
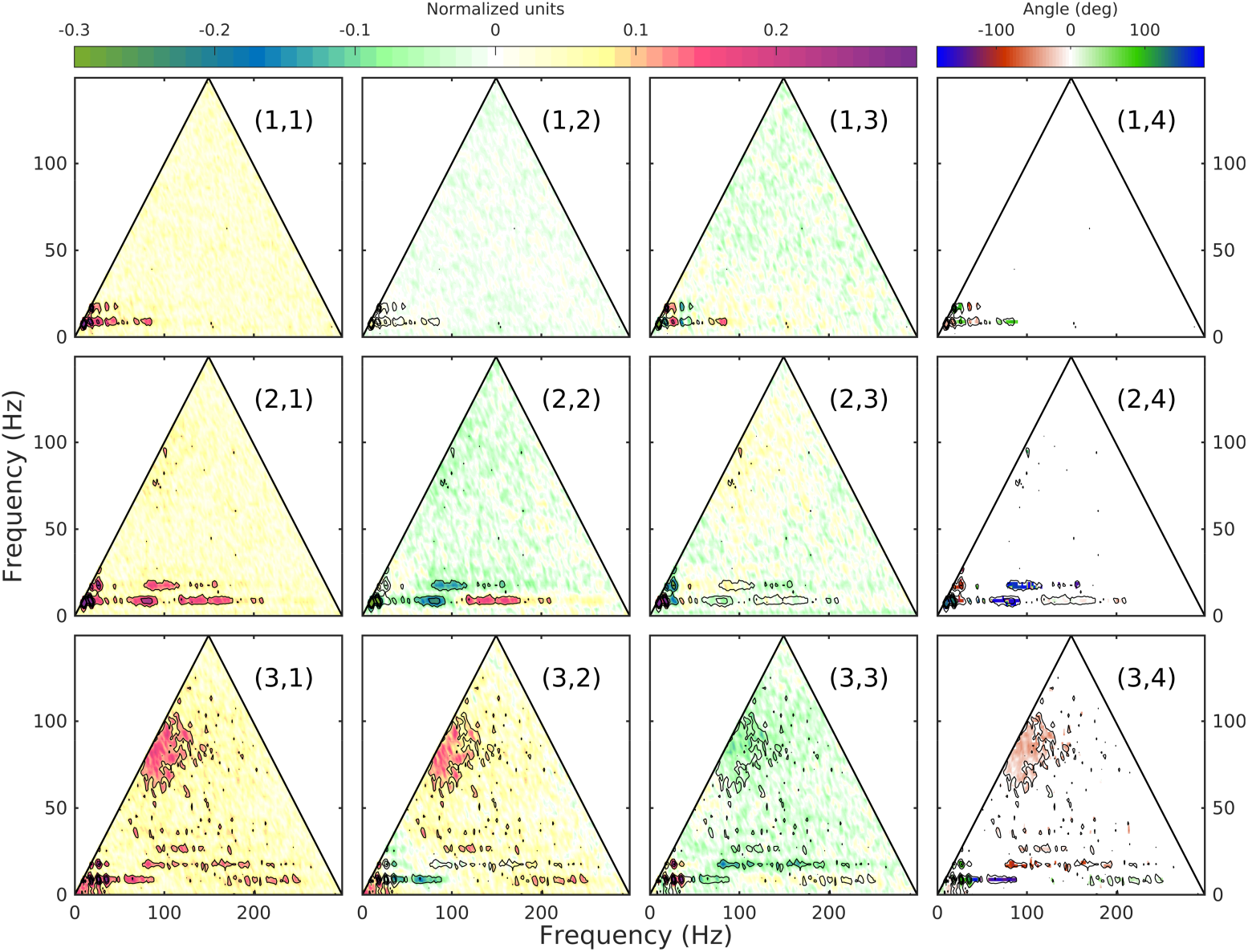
Bispectra of LFP recordings corresponding to high rat speed, *v* > 40 cm/s. Columns: 1) bicoherence (normalized modulus of bispectrum); 2) skewness (normalized real part of bispectrum); 3) asymmetry (normalized imaginary part of bispectrum); 4) biphase (argument of bispectrum). Rows: 1) CA1; 2) LM; 3) DG. See text for an interpretation. CA1 bispectrum shows only phase coupling between theta and its harmonics, suggesting a very weak theta-gamma interaction and cascade. Both LM and DG regions exhibit theta-gamma coupling, indicative of stronger cascade in the inertial gamma range. Remarkably, theta-epsilon coupling can be detected in LM, despite low epsilon power (the epsilon is indistinguishable from the dissipation range). In DG the inertial range cascade extends into the epsilon band. Epsilon is not coupled to theta but seems to be driven by gamma through a much more effective mechanism of second-harmonic excitation with positive skewness, negative asymmetry, and 90 deg. biphase, panels (3,2-4). Gamma contributes negative skewness to the LFP trace, epsilon has a positive contribution, panels (2, 2-3).

**Figure 6:**
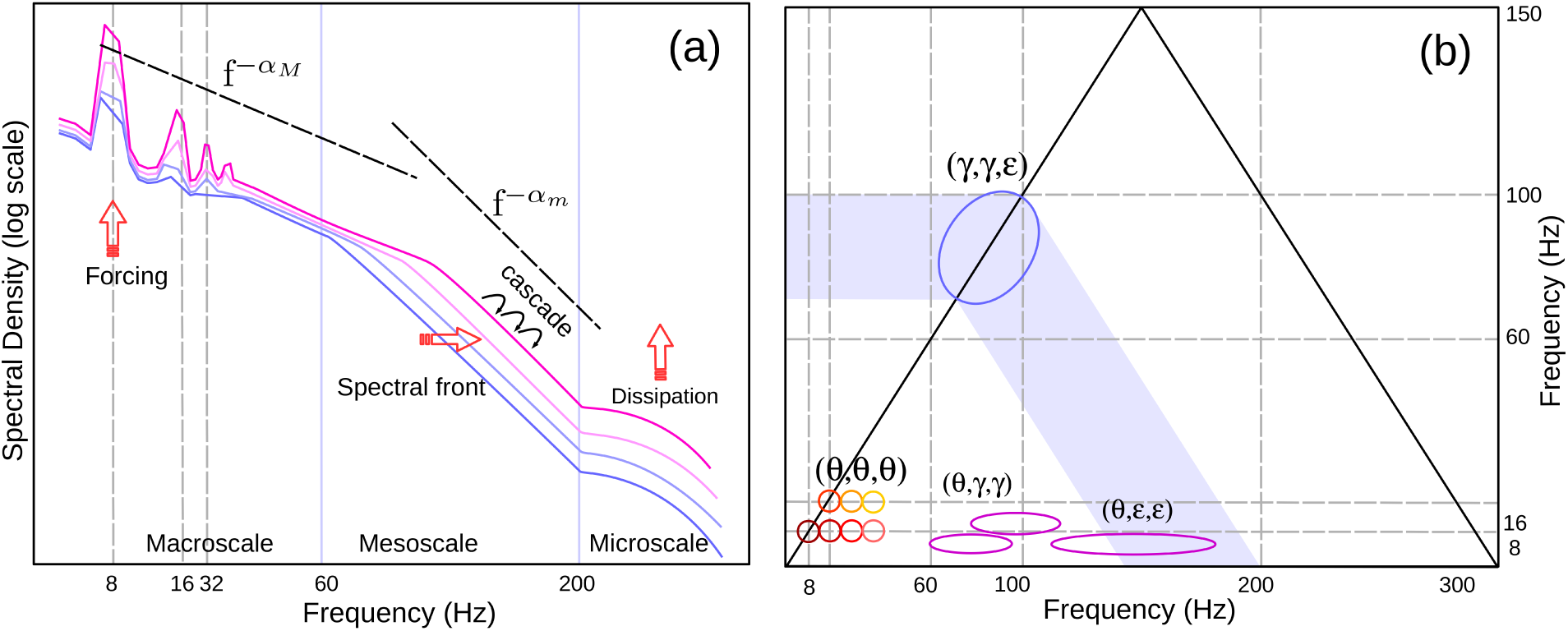
A schematic of main features of hippocampal mesoscale turbulence aggregating information from all three regions investigated, CA1, LM, and DG. See text for an discussion.

## Discussion

Based on recent observations^6, 7^ of wave propagation in the hippocampus, we hypothesize that hippocampal activity (and more general, activity on foliations) can be described as a multi-scale wave field whose dynamics are weakly turbulent. We call this “mesoscale brain turbulence”, and propose to use the theory of weak turbulence to study its physics. The term turbulence is used here in the modern sense pioneered by Zakharov^9^: a branch of statistical mechanics that studies nonlinear systems spanning multiple, weakly-interacting scales. The central concept of the turbulence theory is the spectral cascade, the exchange of energy between scales that occurs in the inertial range of a turbulent system. To our knowledge, brain activity has not been examined from this perspective, and although dynamical models exist^32,33^, the details of wave propagation processes in the brain are not well understood.

The turbulence formalism provides a straightforward dynamical description of neural activity. We illustrate it here by examining the turbulence interpretation of hippocampal statistics. In the context of turbulence, the evolution of hippocampal spectra and bispectra follows a simple scenario, summarized in Fig. 6. Based on position of crossover points, where the spectral slope changes, hippocampal spectra separate naturally into macroscale (global-brain oscillations), mesoscale (local lattice oscillations), and microscale (high-frequency oscillations mainly related to single neuron activity and volleys between neuron pairs). Regardless of their nature in other parts of the brain, on the hippocampal lattice, all oscillations in the macro- and mesoscale range are assumed to be interacting waves. The theta band defines the spectral input window. The microscale identifies with the dissipation range. The mesoscale range (local gamma and epsilon oscillations) is inertial. As rat activity increases, theta amplitude increases and theta harmonics are generated. In the hippocampus, wave interaction mediates the release of theta energy into the mesoscale domain at the low gamma frequencies. The theta-gamma interaction (cascade) is evidenced by the appearance of bispectral peaks of theta-gamma coupling at high speeds (Fig. 6b). Gamma growth with increased rat speed results in a front-like KZ spectrum with a slope *α* ≈ 4.5 that shifts toward higher frequencies, leaving in its wake a macroscale KZ spectrum 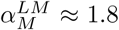, and 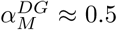, Fig. 6a). In the DG region, the lower gamma energy further cascades into the epsilon domain (a broad bispectral peak of gamma-epsilon coupling appears, Fig. 6c).

For neuroscience, the turbulence promises quantitative insight into hippocampal activity as a function of energy into the network. In the hippocampus, the energy pumped continuously at theta frequency has two effects: it induces a nonlinear deformation of the local theta wave (similar to the forcing in Fig. 1a), and generates a spectral cascade that excites/supports collective activity at the neighboring resonant scales^34,35^. The lower power in the theta band shows that the pyramidal cell layer of CA1 receives the least amount of energy compared to DG and LM, which receive nearly equivalent input. Intuitively, this speaks to the distinction between from fibration to foliations: synapses from external region terminate preferentially within the LM and DG, while the CA1 pyramidal layer region is dominated by local inputs. The CA1 cascade does not appear to develop a KZ spectrum in the gamma range, suggesting of a more dissipative/output-driven function possibly related to relaying activity downstream. The gamma power in the LM is driven via the entorhinal input, paced at the frequency of incoming EPSPs as well as modulated by local interneurnal connections ^27,36^. The epsilon coupling to theta in this region, although not prominent in the power spectra, is a consequence of back-propagating physiology as part of dissipation. The DG epsilon differs from LM in two significant ways: 1) it is more energetic than LM epsilon by an order of magnitude, and 2) it is coupled to gamma as a second-harmonic oscillation. Given the high degree of interneuron activity in the DG, this coupling suggests strong nonlinear interaction between populations of different cell types, most-likely interacting inhibitory networks in order to shape excitatory population firing.Action potential is structured on these resonating interactions, leading to the epsilon-theta coupling seen in the bicoherence plots. This is the clearest evidence of a turbulent cascade leading to a weak coupling of oscillations from the slow theta to fast epsilon

A persistent question is the origin of power-law spectra in the brain^13^. There are several explanations for the occurrence of the *f*^*−α*^ spectrum. The simplest is the fractal noise hypothesis, which describes the brain as directed connections that transmit pulses with a certain life-time distribution (in fact this model only establishes a scale-time duality of self similarity). Billed as a universal model, criticality addresses the question of the origin of the life-time distribution. The mesoscale-turbulence hypothesis presents an alternative explanation for the peculiar, power-law spectra observed on brain foliations. A fundamental result of weak turbulence is the prediction stationary Kolmogorov-Zakharov, power-law spectra, whose exponents are uniquely determined by the physics of the system: the homogeneity of the interaction coefficients, dimensionality of the space, and properties of dispersion relation. Weak turbulence is a powerful formalism that charts a clear path to universal laws governing the stochastic evolution of mesoscale brain activity, and can serve as the foundation for the development quantitative, verifiable models.

## METHODS SUMMARY

The development of this theoretical model utilized biological data from a single rat `(maurer-r539)`, but the spectral characteristics are consistent across other rats (data not shown) and with published studies. The scope of this discussion is the applicability of turbulence principles to brain activity, buttressed by biological data. An in-depth analysis, including statistics across a rat population, is forthcoming and will be published elsewhere.

The rat in the present manuscript was a 4-9 months old Fisher344-Brown Norway Rats. Upon arrival to the University of Florida, the rat was allowed to acclimate to the colony room for one week. After which, the rat slowly trained to traverse a circular track for food reward while their body weight was slowly reduced to 85% to their arrival baseline. Once the rat reliably performed a shuttle task, performing more than one lap per minute, it was implanted with a custom single shank silicon probe from NeuroNexus (Ann Arbor, MI). This probe was designed such that thirty two recording sites, each with a recording area of 177 *µ*m, were spaced 60 um apart allowing incremental recording across the hippocampal lamina.

Following recovery from surgery, the rat was retrained to traverse multiple mazes. For the purposes of this study, data from the circle track maze (outer diameter: 115 cm, inner diameter: 88 cm) was analyzed. The rat was trained to run unidirectionally, receiving food reward at a single location. Local-field potential recordings and video images were recorded synchronously using a Tucker-Davis Neurophysiology System (20 kHz, and 30 frames/s, spatial resolution < 0.5-cm/pixel). All procedures were in accordance with the NIH Guide for the Care and Use of Laboratory Animals and approved by the Institutional Animal Care and Use Committee at the University of Florida.

Details of analysis methods, further supportive data, as well as a brief summary of the main turbulence principles, are given in the Supplementary Material.

## Acknowledgements

This research was supported by National Institute of Mental Health, grant 1R01MH109548-01A1 and National Iinstitute of Aging, grant R01AG055544.

## Author Contribution

A.S. developed the theoretical framework for the application of the turbulence formalism to brain activity, and part of the numerical methods used. Y.Q. developed part of the numerical methods, and performed the numerical analysis. J.P.K managed data collection. A.M. provided the impetus for the research; designed, organized, and supervised the data collection; and provided the guidance and advice regarding all things neuroscience. A.S. wrote the manuscript with input from all authors.

## Competing Interests The authors declare that they have no competing financial interests

